# Brain signaling becomes less integrated and more segregated with age

**DOI:** 10.1101/2023.11.17.567376

**Authors:** Rostam M Razban, Botond B Antal, Ken A Dill, Lilianne R Mujica-Parodi

## Abstract

The integration-segregation framework is a popular first step to understand brain dynamics because it simplifies brain dynamics into two states based on global vs. local signaling patterns. However, there is no consensus for how to best define what the two states look like. Here, we map integration and segregation to order and disorder states from the Ising model in physics to calculate state probabilities, *P*_int_ and *P*_seg_, from functional MRI data. We find that integration/segregation decreases/increases with age across three databases, and changes are consistent with weakened connection strength among regions rather than topological connectivity based on structural and diffusion MRI data.

**AUTHOR SUMMARY:** The integration-segregation framework succinctly captures the tradeoff brains face between seamless function (more integration) in light of energetic constrains (more segregation). Despite its ubiquitous use in the field, there is no consensus on its definition with various graph theoretical properties being proposed. Here, we define the two states based on the underlying mechanism of neuronal coupling strength to provide a physical foundation for the framework. We find that younger adults’ brains are close to perfectly balancing between integration and segregation, while older adults’ brains veer off towards random signaling.

## INTRODUCTION

Aging is the number one risk factor for almost all neurodegenerative diseases (Kennedy et al., 2014). For every 5 years after the age of 65, the probability of acquiring Alzheimer’s disease doubles (Bermejo-Pareja et al., 2008). An influential conceptual approach to begin making sense of brain dynamics frames it in terms of a balance between **integrated** and **segregated** network **states** (Deco, Tononi, Boly, & Kringelbach, 2015; Friston, 2009; Sporns, 2010, 2013; Tononi, Sporns, & Edelman, 1994; Wig, 2017). On one hand, the brain faces functional pressure to have as many regions directly connected for quick communication. On the other hand, the brain is constrained to minimize metabolic energy consumption because it consumes ten-times more of the body’s energy than expected by mass (Raichle, 2006). Tuning the balance between extensive global signaling, referred to as integration, and limited local signaling, referred to as segregation, optimally compromises between functional and energetic constraints (Bullmore & Sporns, 2012; Cohen & D’Esposito, 2016; Manza et al., 2020; Wang et al., 2021). Although these constraints remain throughout life, aging disrupts their balance.

Previous research found mixed aging results, depending on the metrics used to measure integration and segregation (Chan, Park, Savalia, Petersen, & Wig, 2014; Chen et al., 2021; Onoda & Yamaguchi, 2013; Zhang et al., 2021). Although most in the literature use the system segregation metric (Chan et al., 2014), no consensus exists surrounding integration. In general, the problem facing the integration-segregation framework is that there is no one way to define the two states. Many graph theoretical metrics could potentially be used (Rubinov & Sporns, 2010) and it is unclear why one should take precedence over the other, particularly when their aging outcomes are mutually inconsistent. There is a need to more fundamentally define integration and segregation to transform it from a proxy to a physical quantity.

Here, we provide a physical foundation for the framework by applying the mean field **Ising model** to treat integration and segregation as physical 2-**phase** systems like magnets and liquids. After demonstrating that the Ising model can capture global brain dynamics as measured by functional MRI once the effective number of nodes is properly set, we proceed to calculate probabilities of being in the integrated or segregated states and find that younger and older brains are bounded by optimal and random signaling, respectively. We then explore diffusion and structural MRI data to ask if the age-related changes in signaling are due to changes in topological network connectivity.

## APPLYING THE ISING MODEL TO FMRI

We model human brain signaling patterns obtained from resting-state functional MRI (fMRI) data sets. As in previous work (Weistuch et al., 2021), we capture those patterns with the Ising model, a widely used theoretical method for expressing macroscale behaviors in terms of interactions among many underlying microscale agents (Dill & Bromberg, 2012). We first transform the continuous fMRI data into a representation as discrete Ising spins via binarization of the data (Figure 1). That is, we reduce the state of the region as either −1 or 1 based on whether fMRI signaling is decreasing or increasing, respectively. Second, we calculate the *synchrony* by summing over all spins in a given time interval and dividing by the total number of spins (Figure 1). Synchronies are collected over the entirety of the scan to obtain a distribution. Based on Ising model theory, the synchrony threshold delineating between integrated and segregated states is set such that *P*_int_ = *P*_seg_ = 1*/*2 at the Ising model’s **critical point** (Methods). *P*_seg_ is the probability that the brain is in the segregated state and is defined as the relative number of time points for which the absolute value of synchrony is less than the synchrony threshold (Figure 1). *P*_int_ is defined as the relative number of time points for which the absolute value of synchrony is greater than the synchrony threshold and trivially relates to *P*_seg_ because *P*_int_ + *P*_seg_ = 1.

**Figure 1.**
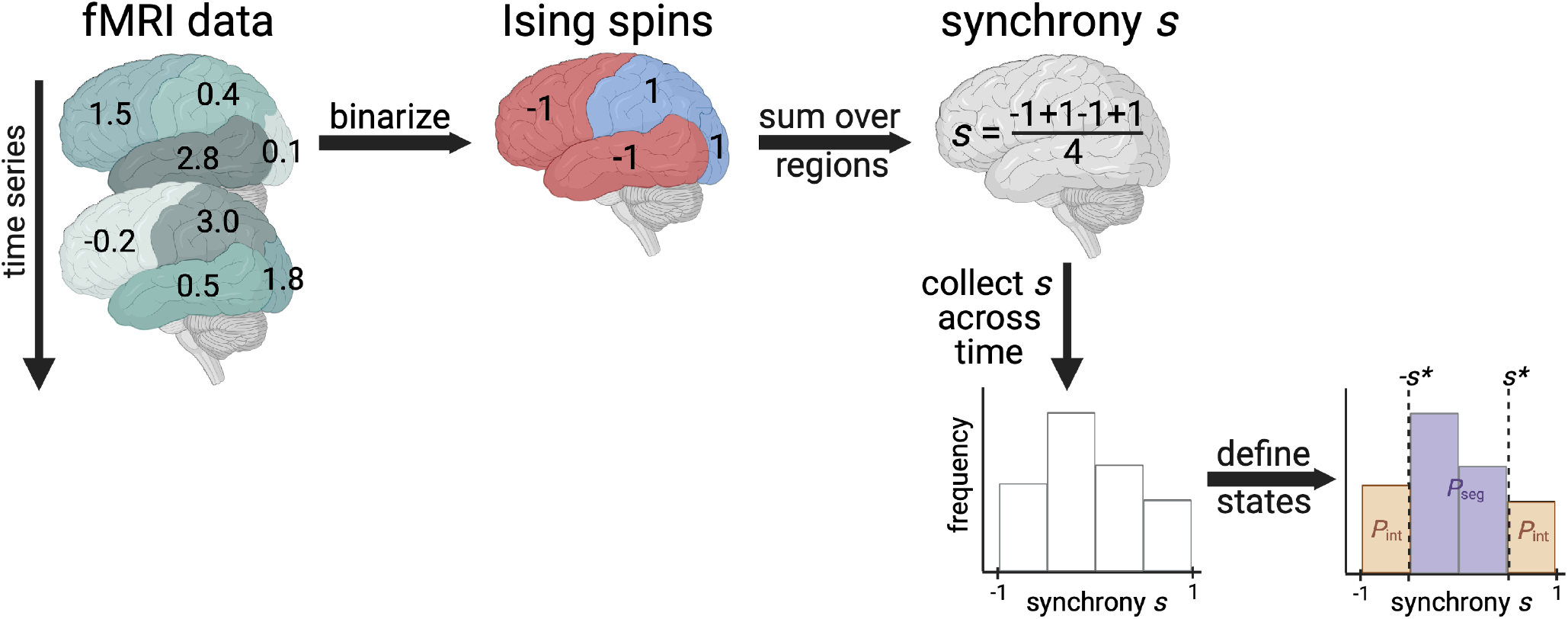
Calculating the probability that the brain exhibits integrated or segregated dynamics (*P*_int_ or *P*_seg_). The schematic demonstrates the procedure for one individual’s fictitious functional MRI scan with 4 brain regions and only two time points shown. First, we binarize data based on nearest neighbor scans in time. If the functional MRI (fMRI) signal increases, a value of 1 is assigned; decreases, -1. Then, we calculate the average spin state of the brain, called synchrony. Finally, we collect synchrony values across the entire time series to create a synchrony distribution. We appropriately set the synchrony threshold based on Ising model theory to delineate between integrated and segregated microstates. Additional details can be found in the Methods. Figure created with Biorender.com.

## RESULTS

### The number of functionally effective brain regions

Before proceeding to calculate *P*_seg_, we first check whether the model can capture the experimental synchrony distributions. A mean field Ising model only considering pairwise interactions has one quantity of interest. The strength of connection *λ* between any two regions corresponds to the degree to which signals between any two brain regions are correlated. However, we find that a naive fit of *λ* based on **maximum entropy** (Dill & Bromberg, 2012; Schneidman, Berry, Segev, & Bialek, 2006; Weistuch et al., 2021) fails to capture the synchrony distribution from fMRI data (Figure 2, orange). To improve upon a standard Ising model approach, here we introduce a hyper-parameter *N*_eff_. Brain atlas parcellations provide *N* brain regions, however, those *N* regions must be identically distributed across time for the Ising model to apply. We find that when setting *N* to a lower value *N*_eff_, fixed for all individuals within a data set, the Ising model accurately captures synchrony distributions (Figure 2). The optimal value of *N*_eff_ = 40 is determined by scanning across *N*_eff_ multiples of 5 to find which best captures the next order moment not fit by our maximum entropy setup across all individuals (Methods, Figure 6). For our particular preprocessing (Methods), we find that *N*_eff_ = 40 for individuals in the Cambridge Center for Ageing and Neuroscience (CamCAN) (Taylor et al., 2017) and the Human Connectome Project Aging (HCP) (Harms et al., 2018). For the UK Biobank (UKB) (Alfaro-Almagro et al., 2018), *N*_eff_ = 30 performs best (Figure 6).

**Figure 2.**
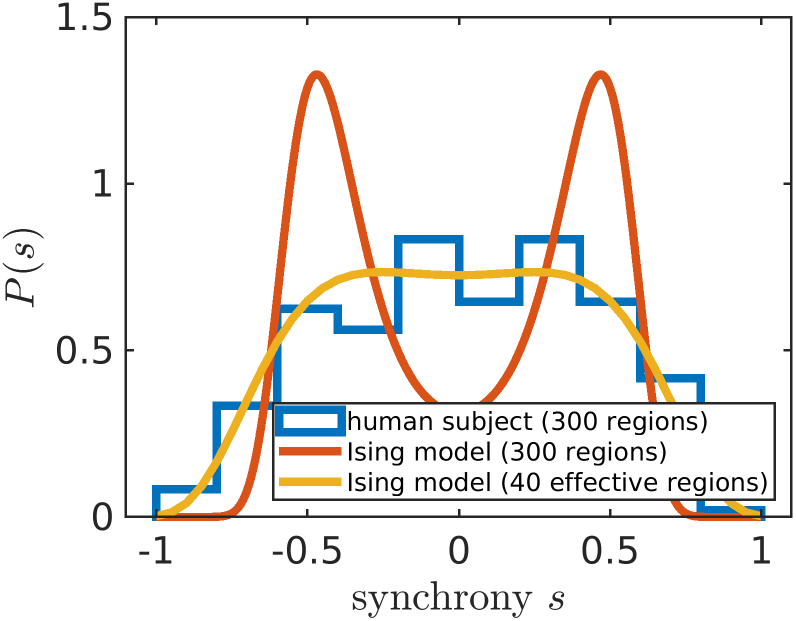
Adjusting the number of brain regions (*N*_eff_) helps capture experiment. The modified Ising model with *N*_eff_ = 40 (yellow line) better captures the synchrony distribution (blue histogram) of an arbitrarily chosen individual in the Cambridge Centre for Ageing and Neuroscience data set (subject id: CC110045). The orange line corresponds to the Ising model with *N* equal to the number of regions in the Seitzman atlas (Seitzman et al., 2020).

Based on identified *N*_eff_ hyper-parameter values, brains act as if they have a few tens of functional units. If different preprocessing decisions are considered, such as atlas resolution, *N*_eff_ values are still within an order of magnitude. At the voxel-level (*N* = 125, 879), we obtain an *N*_eff_ value of 65 for CamCAN and 125 for HCP using the same procedure as for the Seitzman atlas (*N* = 300) considered in the previous paragraph (Figure S2). Future work will pinpoint how *N*_eff_ depends on preprocessing to enable a future study creating a physics-based parcellation of the brain.

We also tried an alternative fitting strategy by fitting *N*_eff_ per individual rather than having the same value for all individuals in a respective data set. We show that individually fitted *N*_eff_ values trivially relate to *λ* as expected by theory (Figure S1). Moreover, individually fitted *N*_eff_ are not found to be related to global differences in anatomical brain connectivity (Figure S3).

### The aging brain becomes functionally more segregated

With an appropriately determined *N*_eff_, we can accurately set the same synchrony threshold *s*^*^ for all individuals within a data set to calculate *P*_seg_. The value of *s*^*^ is set such that at the Ising model’s critical point in connection strength *λ, P*_seg_ equals to 1*/*2 for the ideal synchrony distribution based on Ising model theory (Methods). This enables *P*_seg_ comparisons across data sets that may have different *N*_eff_ values. For CamCAN and HCP, *s*^*^ = 0.33 because *N*_eff_ = 40 for both data sets. For UKB, *s*^*^ = 0.36 (Table S1).

Across the three publicly available data sets, we find that the balance shifts towards more segregation at older ages (Figure 3). Note that if we plotted *P*_int_ rather than *P*_seg_, Figure 3 would be horizontally flipped, where *P*_int_ goes from high to low values as a function of increasing age because *P*_seg_ + *P*_int_ = 1. There is large variation among subjects (Figure S4). However, the correlation between age and *P*_seg_ is significant with the largest coefficient being 0.40 for CamCAN, while the lowest being 0.08 for UKB. Discrepancies in study designs may explain correlation magnitude differences: CamCAN and HCP are designed to study healthy aging (Bookheimer et al., 2019; Shafto et al., 2014), while the goal of UKB is to identify early biomarkers for brain diseases (Sudlow et al., 2015).

**Figure 3.**
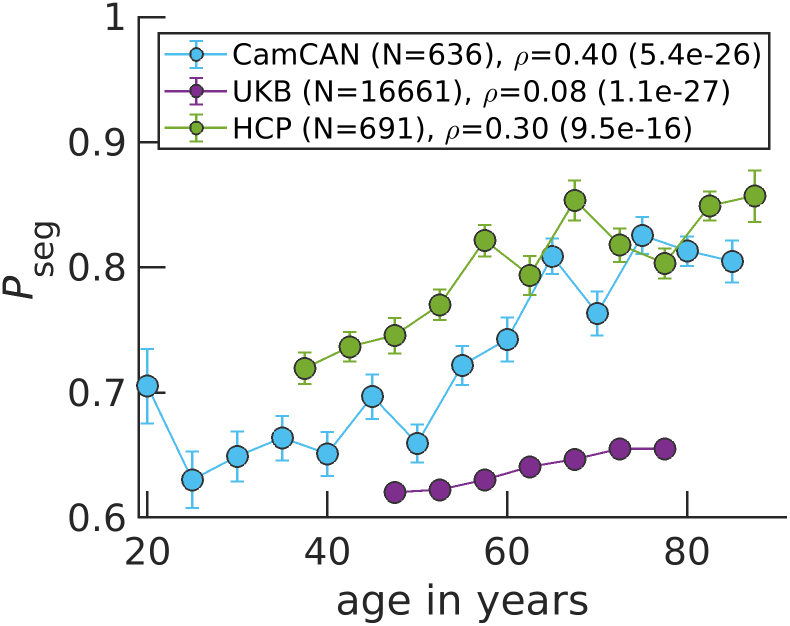
*P*_seg_ rises in aging brains across three data sets. Data points correspond to medians, while error bars correspond to standard errors for bins of 5 years. The variable *ρ* corresponds to the Spearman correlation coefficient between age and *P*_seg_ calculated over all N individuals, with the p-value in parenthesis.

To better highlight how *P*_seg_ changes across CamCAN’s large age range, we present violin plots for younger, middle age and older individuals’ *P*_seg_ (Figure S5). We also investigate how *P*_seg_ varies across time for a given individual. In Figure S6, we show that the per individual *P*_seg_ standard deviations decrease across age for CamCAN and HCP individuals. Finally, we perform a multiple linear regression with sex and handedness as additional covariates and show that age still strongly explains increasing segregation (Table S2, S3 and S4; Figures S7, S8 and S9).

Informed by the Ising model, increases in segregation result from network reorganization to more local signaling because of weakened connection strength between regions. Interestingly, younger individuals exhibit segregation behavior closer to the Ising model’s critical point of connection strength (Figure S10). At the critical point, we define *P*_seg_ = 1*/*2 (Methods) and find experimental *P*_seg_ values closer to 1*/*2 for younger individuals (Figure 3). Older individuals on the other hand, approach *P*_seg_ = 1 on average. This limit corresponds to functionally uncoupled brain regions that are randomly activating. Our results support the critical brain hypothesis that healthy brains operate near a critical point (Beggs, 2022; Beggs & Plenz, 2003; Ponce-Alvarez, Kringelbach, & Deco, 2023; Tagliazucchi, Balenzuela, Fraiman, & Chialvo, 2012) and implicate aging as pushing brain dynamics further away from criticality.

### Increasing segregation is not related to structural degradation

In the previous subsection, we discussed the disruption of the integration and segregation balance from the perspective of phase transitions in physics. Here, we explore the physiological mechanism underlying increasing segregation in the aging brain. We consecutively simulate the Ising model on a hypothetically degrading brain structure and show that random removal of edges yields qualitatively similar results to those of fMRI (Figure 4). Note that Figure 4 is horizontally flipped from those of *P*_seg_ (Figure 3) because average degree (relative number of edges) is on the x-axis. It is presumed that edges are lost as age increases. In Figure 4, edges are lost linearly in time, however, more complicated monotonic functions can be employed to yield a quantitative match with experimental data in Figure 3. We can also capture variability among individuals by assuming connection strengths within an individual are drawn from a distribution, rather than all being equal (Figure S11). In the supplement, we also demonstrate that similar qualitative trends are obtained when starting with other individuals’ structures, regardless of their age (Figure S12).

**Figure 4.**
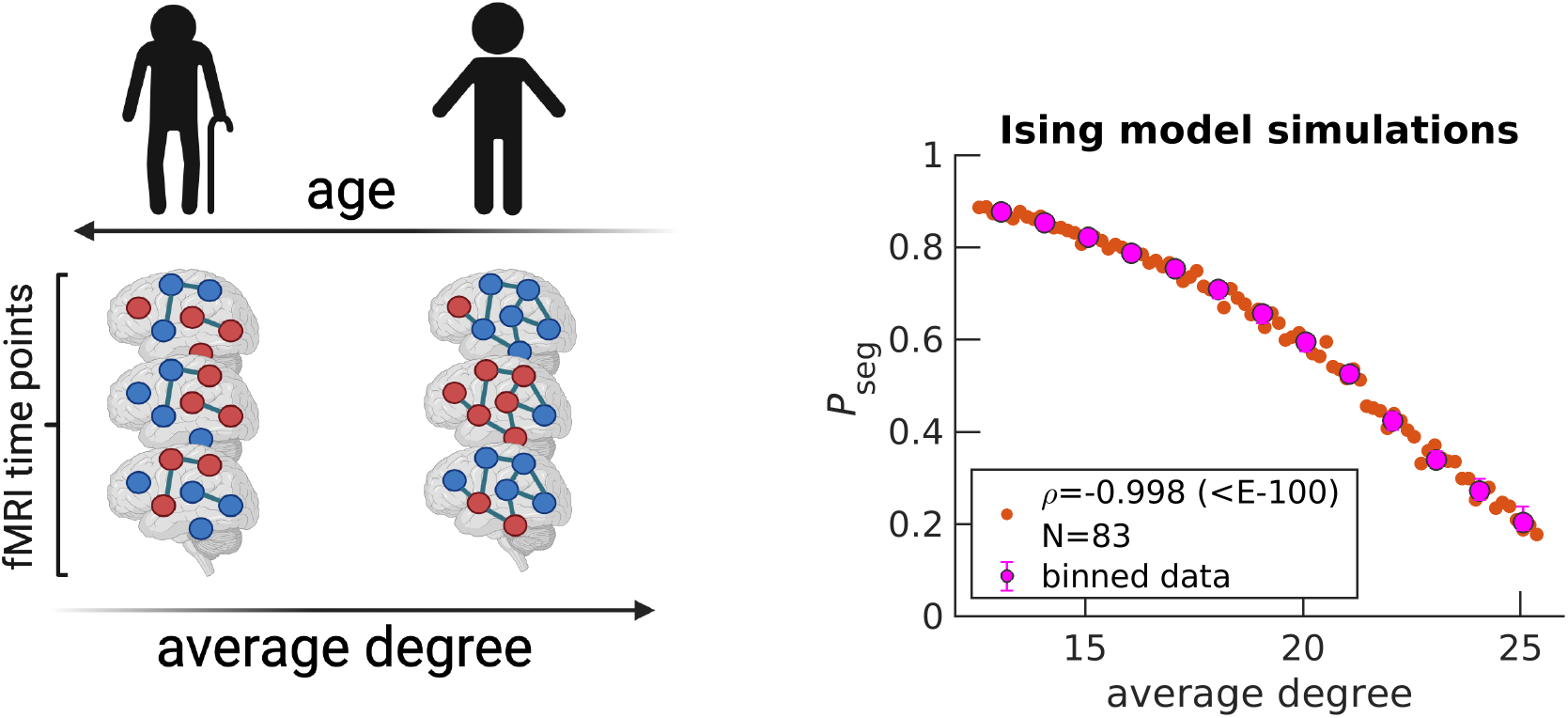
Simulating the random removal of edges results in *P*_seg_ increases. Five edges are randomly removed from a starting diffusion MRI structure (arbitrarily chosen UK Biobank individual, subject ID: 6025360, 51 years old), under the Harvard-Oxford atlas (64 regions). An Ising system is simulated with *N*_eff_ = *N* = 64 for the corresponding diffusion MRI structure. Spin states, denoted by dark blue and red node colors in the schematic, are recorded across 2500 time steps to calculate *P*_seg_. Then, the entire procedure is repeated for the updated structure after edge removal, for a total of 83 times (Methods). Orange data points on the right plot correspond to individual Ising systems, where N reflects the total number. The variable *ρ* corresponds to the Spearman correlation coefficient calculated over all orange data points between average degree and *P*_seg_, with the p-value in parenthesis. Magenta data points correspond to medians, while error bars correspond to upper and lower quartiles for bin sizes of one degree. The schematic on the left is created with Biorender.com.

We now begin to investigate possible mechanisms of connection degradation. First, we find that our simulation is agnostic to the detailed mechanism of connection degeneration because connection strength is essentially modulated by the probability that a given edge exists (Figure S13). In other words, the simulation cannot inform whether connections are degraded based on some targeted property. Thus, we turn to structural MRI and diffusion MRI data from UKB to investigate possible properties being degraded with age. In Figure S15, we confirm that **white matter** volume decreases as a function of adult age, as previously reported (Bethlehem et al., 2022; Lawrence et al., 2021; Lebel et al., 2012). However, this decrease does not correspond to a loss of anatomical connections because we find that neither average degree, average tract length nor average tract density monotonically decrease with age when analyzing diffusion MRI scans using the Q-Ball method (Figure S16). This seems to contradict previous findings which report decreases (Betzel et al., 2014; Lim, Han, Uhlhaas, & Kaiser, 2015). However, previous results employed the more simple diffusion tension imaging (DTI) method which is known to be less accurate at performing tractography (Garyfallidis et al., 2014; Jones, Knösche, & Turner, 2013; Rokem et al., 2015). When rerunning our analysis for DTI, we can reproduce previously reported tract properties’ anticorrelations with age (Figure S16). We also investigate a graph property that captures polysynaptic connectivity called communicability (Andreotti et al., 2014; Estrada & Hatano, 2008; Seguin, Sporns, & Zalesky, 2023) and find that it also does not decrease age when using Q-Ball derived tract density (Figure S17).

We propose that observed white matter volume reduction (Figure S15) and brain dynamics change corresponds to less myelin covering axons as function of age. Despite rejecting anatomical connections as a possible mechanism in the previous paragraph, it remains inconclusive whether myelin underlies trends because we are not aware of such data being publicly available. Although axons are still physically present, myelin coverage disruption causes regions to no longer be functionally connected because signals do not arrive on time. Previously reported results from Myelin Water Imaging confirm reduction in myelin at advanced ages (Arshad, Stanley, & Raz, 2016; Buyanova & Arsalidou, 2021). We also investigated whether degraded functional connections are likely to be longer than average with age, as previously reported for certain brain regions (Tomasi & Volkow, 2012). Although we indeed find that the average correlation of the 25% longest connections is slightly more strongly anticorrelated with age compared to the average correlation of the 25% shortest connections for CamCAN (Figure S18, left), we find the opposite trend for HCP (Figure S18, right). Thus, myelin reduction does not seem to have a stronger impact on longer connections and conclude that the loss of functional connections happens randomly with respect to length at the brain-wide scale.

## DISCUSSION

We apply the mean field Ising model to physically quantify integration and segregation at the emergent scale of the whole brain. From resting-state fMRI scans across three publicly available data sets, we find that brain dynamics steadily becomes more segregated with age. Physically, aging leads to brain dynamics moving further away from its optimal balance at the critical point. Physiologically, analyses of white matter properties point to random functional connection losses due to myelin degeneration as the possible culprit for more segregated dynamics. This expands upon our previous work finding metabolic dysfunction to underly brain aging (Weistuch et al., 2021), hinting that myelin may be especially vulnerable to energy imbalances.

The Ising model and integration-segregation frameworks are considered as the simplest approaches to capture dynamics in their respective fields. Thus, it is fitting to map segregated and integrated states in neuroscience to disordered and ordered Ising model phases in physics, respectively. One general challenge in applying graph theory to MRI-level data is identifying what constitutes a node (DeFelipe, 2010; Lacy & Robinson, 2020; Seung, 2012; Sporns, 2010; Wig, Schlaggar, & Petersen, 2011; Yeo & Eickhoff, 2016). We identify the best number of effective brain regions *N*_eff_ such that the Ising model accurately captures individuals’ synchrony distributions across the corresponding data set, improving upon our original application of the Ising model which lacked the *N*_eff_ hyper-parameter (Weistuch et al., 2021). Future work will utilize *N*_eff_ calculations to guide the creation of a parcellation in which brain regions are constrained to be physically independent based on their collective functional activity.

The field is inundated with integration and segregation metrics that have different aging trends. We go beyond heuristic definitions, such as one that we previously proposed based on matrix decomposition (Weistuch et al., 2021), by self-consistently defining the two states within the Ising model framework.

This makes our metric mechanistically based on the connection strength between regions and further stands out because *P*_seg_ and *P*_int_ are naturally at the emergent scale of the brain. We do not calculate a local property and then average over nodes to yield a brain-wide value ((Wang et al., 2021)’s metric also has this advantage). In addition, *P*_seg_ and *P*_int_ are directly related because *P*_seg_ + *P*_int_ = 1. Most integration and segregation metrics (Chan et al., 2014; Rubinov & Sporns, 2010; Tononi et al., 1994; Wang et al., 2021) are not defined to be anti-correlated. This could be advantageous because greater complexity can be captured (Sporns, 2010).

Taken together, it is not surprising that *P*_seg_ and *P*_int_ results are not consistent with some previous aging reports. For example, a property called system segregation, defined as the difference between inter- and intra-correlations among modules, was found to decrease with age (Chan et al., 2014). Although most report that segregation decreases with age, regardless of the specific metric (Chan et al., 2014; Damoiseaux, 2017; King et al., 2018; Zhang et al., 2021) (see (Chen et al., 2021) for an exception), integration trends are less clear. Global efficiency, taken from graph theory, was found to increase with age (Chan et al., 2014; Yao et al., 2019); however, others found different integration metrics decreasing with age (Chong et al., 2019; Oschmann, Gawryluk, & Initiative, 2020; Zhang et al., 2021), consistent with results reported here.

The utility of the integration-segregation framework lies in its simplicity. However, its simplicity has led to various heuristic definitions that have qualitatively different aging trends. By physically defining integration and segregation based on connection strength between regions, we provide an interpretable foundation for more detailed studies going beyond the two-state approximation to investigate brain dynamics.

## METHODS

### fMRI preprocessing

We access three publicly available resting-state functional MRI data sets: Cambridge Centre for Ageing and Neuroscience (CamCAN) (Taylor et al., 2017), UK Biobank (UKB) (Alfaro-Almagro et al., 2018), and Human Connectome Project (HCP) (Harms et al., 2018). Acquisition details such as field strength and repetition time can be found in Table S5. Demographic details can be found in Table S6.

UKB and HCP fMRI data are accessed in preprocessed form (for details see (Alfaro-Almagro et al., 2018) and (Glasser et al., 2018, 2013), respectively). We preprocessed CamCAN data as done in our previous work (Weistuch et al., 2021). For all three data sets, the cleaned, voxel space time series are band-pass filtered to only include neuronal frequencies (0.01 to 0.1 Hz) and smoothed at a full width at half maximum of 5 mm. Finally, we parcellate into 300 regions of interest according to the Seitzman atlas (Seitzman et al., 2020). For our voxel-wide analysis presented in the Supporting Information, we do not perform parcellation and just consider gray mater voxels by masking.

Applying the Ising model requires the data to only take two possible values: −1 or 1. After performing the preprocessing outlined in the previous paragraph, we binarize the continuous signal for a given region based on the sign of the slope of subsequent time points (Weistuch et al., 2021). We previously showed that such binarization still yields similar functional connectivities as that of the continuous data (Weistuch et al., 2021).

Finally, we only consider brain scans that have the same number of measurements as the predominant number of individuals in the respective data set (Table S5). If the fitted connection strength parameter *λ* is less than 0, reflecting a nonphysical value, we do not include that individual’s brain scan in our analysis. In the HCP data set, we excluded individuals aged 90 years or older because their exact age, considered protected health information, is not available.

### Identifying the N_eff_ hyper-parameter

In Figure 2, our maximum entropy fit (orange line) fails to qualitatively capture the synchrony distribution for an arbitrary individual. To rescue the fit, we replace *N* with *N*_eff_ (Equation S1). In the right plot of Figure 5, we demonstrate that a mean field Ising model with *N*_eff_ = 40 accurately captures the fourth moment of synchrony ⟨*s*^4^⟩ across all individuals in CamCAN preprocessed under the Seitzman atlas. Note that *N*_eff_ is not a parameter like Λ; rather it is a hyper-parameter because it takes the same value across all individuals within the data set. *N*_eff_ is necessary because the Ising model systematically underestimates ⟨*s*^4^⟩ when Λ *>* 0 (left plot of Figure 5). Note that Λ corresponds to rescaling *λ* such that Λ = 0 is at the critical point (Equation S13).

**Figure 5.**
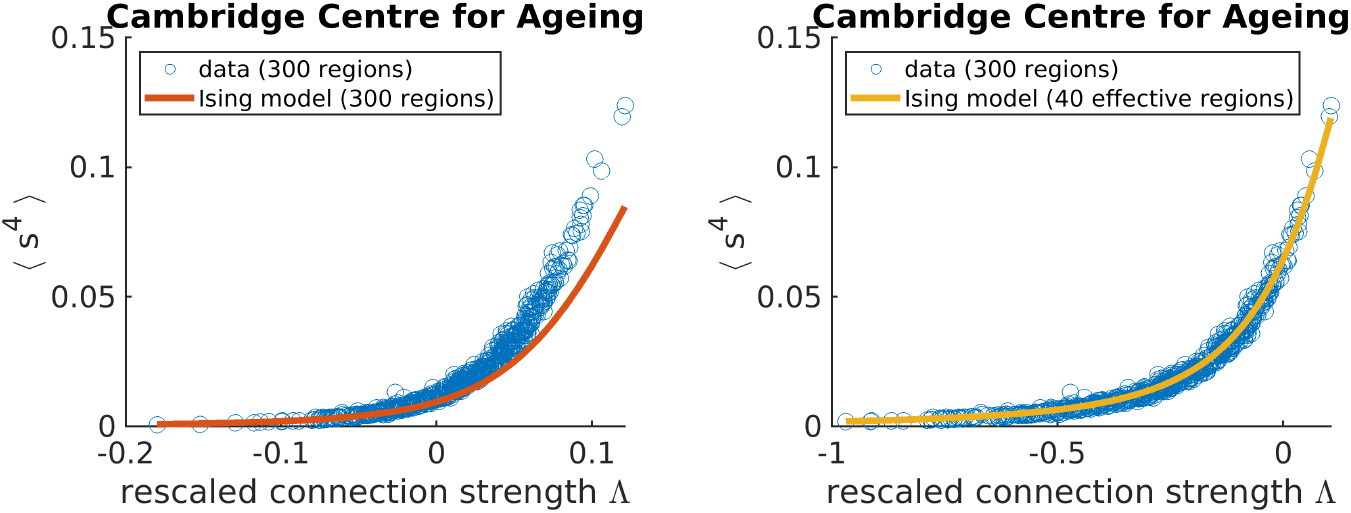
Adjusting the effective number of brain regions (*N*_eff_) helps capture synchrony distributions’ variances across individuals in the Cambridge Centre for Ageing data set. Each data point corresponds to an individual.

**Figure 6.**
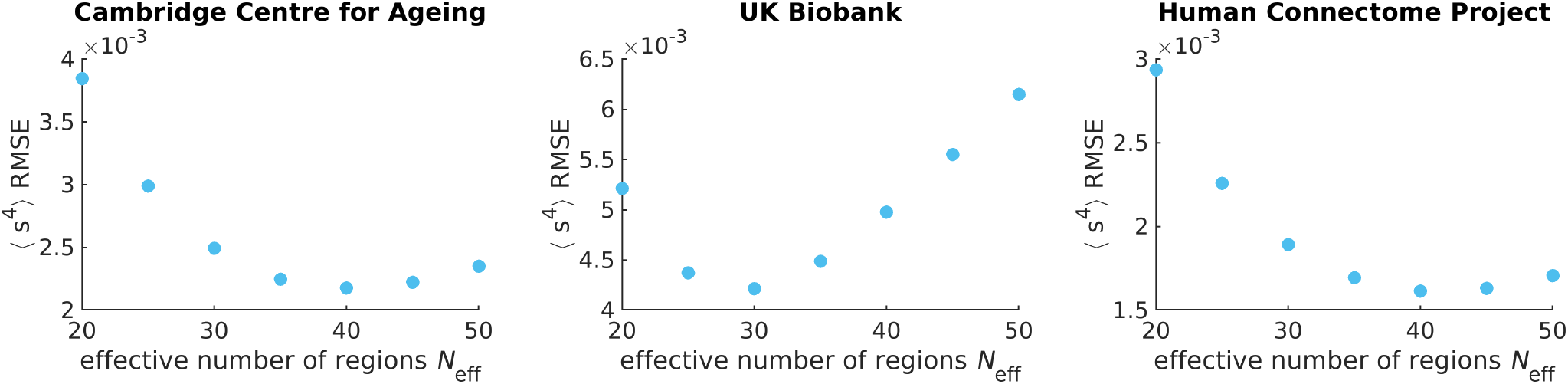
The effective number of regions *N*_eff_ is identified by minimizing the root mean square error (RMSE) of the fourth moment of synchrony between theory and experiment across all individuals. Each data point corresponds to the sum over all individuals’ RMSEs in the respective data set. Note that the y-axis should be scaled by 10^−3^.

To identify *N*_eff_ = 40 as the best value, we perform a parameter scan over multiples of 5 and identify the *N*_eff_ at which the root mean square error (RMSE) between ⟨*s*^4^⟩_exp_ and ⟨*s*^4^⟩_model_ is minimized (Figure 6). We choose the fourth moment because it is the next order moment that our maximum entropy fit does not constrain. It is not the third moment because the distribution is assumed to be even as indicated by our prior (Equation S1).

### Calculating P_seg_

The probability of the brain network being in the segregated state is the sum over all microstates corresponding to the segregated state.

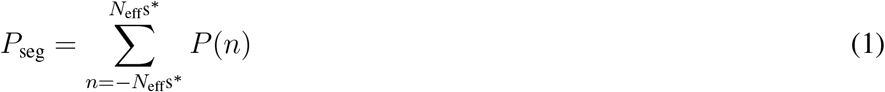

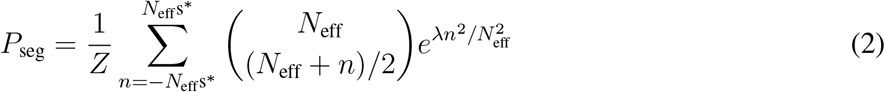

In the second line, the mean field Ising model’s *P* (*n*) is inserted (Equation S2). *Z* corresponds to the partition function and ensures that *P* (*n*) is normalized. The constant *s*^*^ is the synchrony threshold for which segregated and integrated microstates are delineated. We set *s*^*^ such that *P*_seg_ = 1*/*2 when Λ = 0 according to theory. More specifically, we numerically calculate *P*_seg_(Λ = 0) for a given *N*_eff_ and extrapolate to find *s*^*^ (Figure S19). Proper calibration ensures that the theory is accurate and enables apples to apples *P*_seg_ comparisons across data sets with different *N*_eff_. The list of *s*^*^ values for the three publicly available data sets studied can be found in Table S1.

### Ising model simulation

We simulate the Ising model on an initial structure informed by diffusion MRI under the Harvard-Oxford atlas (Makris et al., 2006) (64 regions) for an arbitrarily chosen UK Biobank individual (subject ID: 6025360). If no edge exists between two regions, then the regions are uncoupled. If an edge does exist, then regions *i* and *j* are coupled and contribute *λ* * *σ*_*i*_ * *σ*_*j*_ to the system’s energy; where *λ* corresponds to the connection strength and *σ* corresponds to the spin state of the corresponding region (−1 or 1). Under the standard notation of the Ising model, *λ* = *J/T*, where *J* corresponds to the coupling constant and *T* is the temperature of the bath. The starting *λ* is set to 34.4, which is above *λ*’s critical point (starting *P*_seg_ ≈ 0.2). By definition, *N*_eff_ = *N* = 64 in the simulations. Based on atlas resolution, simulating the Harvard-Oxford atlas provides an *N*_eff_ similar to those found for the experimental data (*N*_eff_ = 40 for CamCAN and HCP; *N*_eff_ = 30 for UKB).

The simulation for a given structure starts by randomly assigning the 64 nodes up or down spins. Then, for each time step, we attempt 10 spin flips 64 times, for a total of 2500 time steps. Spin flips are accepted according to the Metropolis-Hastings algorithm (Metropolis, Rosenbluth, Rosenbluth, Teller, & Teller, 1953). The exact number of spin flip attempts or total time points does not matter, as long as equilibrium is reached. For example, we find that for *λ* values larger than those presented in the text, synchrony distributions become asymmetric and exhibit only one of the two peaks corresponding to the integrated state because of the high kinetic barrier of going from all down spins to all up spins.

Although the starting structure is informed by diffusion MRI, resulting structures after computational edge removals are based on the posited removal strategy. Edges informed by dMRI are undirected and removal maintains undirectedness. Effectively two times as many edges are removed because both forward and backward edges are concurrently eliminated. In Figure S14, we demonstrate how synchrony distributions change as edges are computationally removed for a UK Biobank individual (subject ID: 6025360), with a starting *λ* = 34.4.

We also investigate other individuals’ structures in the UK Biobank to test the robustness of our qualitative results. We arbitrarily chose the following six individuals to widely sample different ages; subject IDs: 6025360 (51y), 4712851 (57y), 3081886 (61y), 1471888 (65y), 4380337 (72y), and 1003054 (74y) (Figure S12). To ensure that the starting *λ* are comparable despite differing in the probability that two regions are connected (*p*_edge_), we set *λ*_0_ = 86.0 for all simulations such that *λ* = *λ*_0_ * *p*_edge_. For example, for subject ID: 6025360, *p*_edge_ = 0.40, thus the starting *λ* = 34.4.

### Diffusion MRI analysis

Diffusion MRI processing to obtain structural information such as tract length and streamline count, which we call tract density, is outlined in our previous work (Razban, Pachter, Dill, & Mujica-Parodi, 2023). Briefly, we take preprocessed dMRI scans from the UK Biobank (Sudlow et al., 2015) and calculate connectivity matrices using the Diffusion Imaging in Python software (Garyfallidis et al., 2014). We input the Talairach atlas (Lancaster et al., 2000) to distinguish between white and gray matter. We perform deterministic tractography and reconstruct the orientation distribution function using Constant Solid Angle (Q-Ball) with a spherical harmonic order of 6 (Aganj et al., 2010). For Figure S16, we also do reconstruction using diffusion tensor imaging (Garyfallidis et al., 2014). To generate the starting structure for Ising model simulations, we input the Harvard-Oxford atlas for tractography because it parcellates the brain into fewer regions, making it more computationally tractable to carry out simulations and closer to *N*_eff_ values found for experimental data.

## Code and data availability

Scripts necessary to reproduce figures and conclusions reached in the text can be found at github.com/rrazban/2state brain. Please refer to the respective publicly available data set to access previously published data (CamCAN, UKB and HCP) (Alfaro-Almagro et al., 2018; Harms et al., 2018; Taylor et al., 2017).

## ACKNOWLEDGMENTS

We thank Ying-Jen Yang, Anthony Chesebro, Charles Kocher, Jonathan Pachter and Lakshman Verma for insightful discussions. The research described in this paper is funded by the White House Brain Research Through Advancing Innovative Technologies (BRAIN) Initiative (NSFNCS-FR 1926781 to L.R.M.-P. and K.A.D.) and the Stony Brook University Laufer Center for Physical and Quantitative Biology (K.A.D.).

Data collection and sharing for this project was provided by the Cambridge Centre for Ageing and Neuroscience (CamCAN). CamCAN funding was provided by the UK Biotechnology and Biological Sciences Research Council (grant number BB/H008217/1), together with support from the UK Medical Research Council and University of Cambridge, UK.

This research has been conducted using the UK Biobank Resource under Application Number 37462. Research reported in this publication was supported by the National Institute On Aging of the National

Institutes of Health under Award Number U01AG052564 and by funds provided by the McDonnell Center for Systems Neuroscience at Washington University in St. Louis. The HCP-Aging 2.0 Release data used in this report came from DOI: 10.15154/1520707. The HCP data repository grows and changes over time. The HCP data used in this report came from NIMH Data Archive DOI:10.15154/1526427.

## SUPPORTING INFORMATION

### Deriving the Ising model with N_eff_

Many have derived the probability distribution of the mean field Ising model, otherwise known as the fully connected or Curie-Weiss Ising model (Friedli & Velenik, 2017; Kochmański, Paszkiewicz, & Wolski, 2013; Weistuch et al., 2021). Here, we demonstrate how to introduce *N*_eff_ in a Maximum Entropy framework. The tricky part is that *N*_eff_ defines the state space over which the probability distribution is summed.

Adding a global pairwise correlation constraint, we obtain the following Lagrangian function *ℒ* over the net displacement of spin states 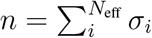, where *σ*_*i*_ can take a value of 1 or −1.

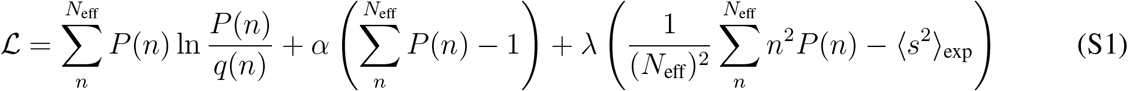

*P* is the probability distribution. *q* corresponds to the prior and is set to the binomial distribution 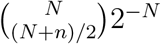, where the binomial coefficient captures the number of ways individual spins can organize for a given *n. α* and *λ* correspond to the Lagrange multipliers that enforce the constraints that the probability distribution is normalized and the mean pairwise correlation equals ⟨*s*^2^⟩, respectively. The variable *s* corresponds to the synchrony, or commonly referred to as the magnetization in ferromagnetic applications, and is limited to vary from −1 to 1 because *n* = *N*_eff_*s*. This is the reason *N* does not have to be the same for ⟨*s*^2^⟩_exp_ and ⟨*s*^2^⟩_model_; ⟨*s*^2^⟩ is always bounded between −1 and 1.

Maximizing the Lagrangian function (Equation S1) with respect to *P*, we obtain the following distribution:

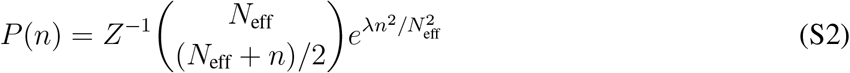

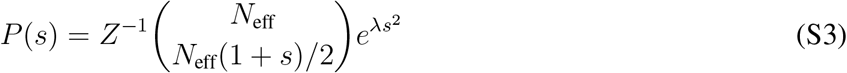

*Z* corresponds to the partition function and ensures that *P* is normalized. The *α* Lagrange multiplier is not present in the final expression because it is subsumed by *Z*.

### Ising model phase transitions

The Landau model is a general formulation to study phase transitions (Dill & Bromberg, 2012; Landau, 1937). It takes the following form,

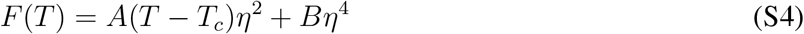

*η* corresponds to the order parameter. *F* is the free energy and can be expressed as the probability for being in microstate *i* by the following relationship *F*_*i*_ = *k*_*b*_*T* ln *P*_*i*_. *T* is the temperature and *T*_*c*_ corresponds to the critical temperature at which a second-order phase transition occurs. *A* and *B* are constants.

Here, we will express the Ising model’s probability distribution (Equation S3) in terms of the Landau formalism (Equation S4) by approximating the binomial coefficient as an exponential to order (*s*^4^). For brevity, we will write *N* to represent *N*_eff_. First, we use Stirling’s approximation to expand the binomial coefficient.

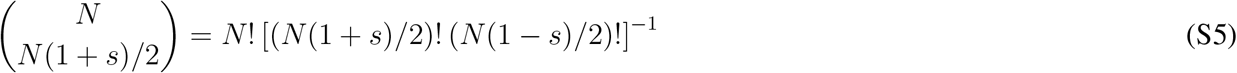

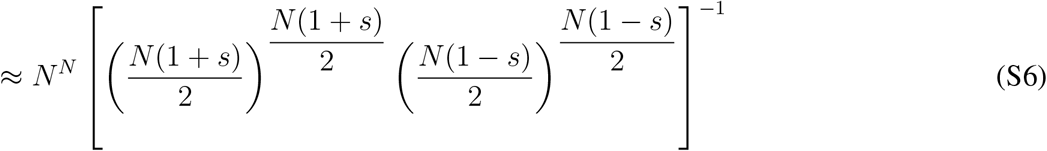

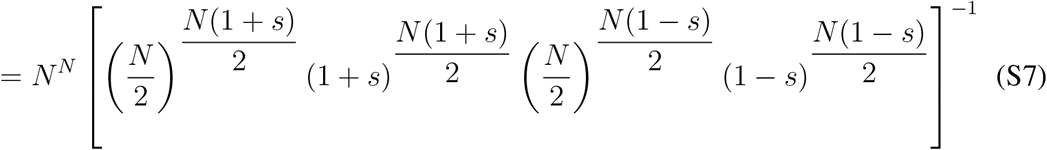

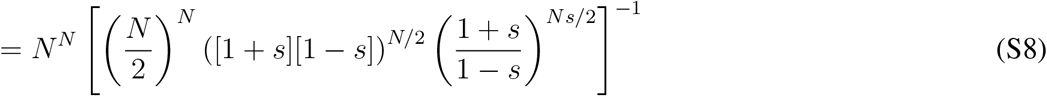

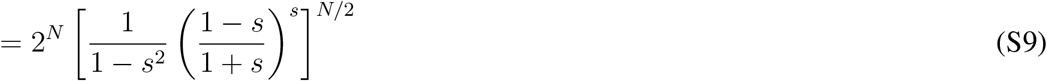

To make further headway, we assume that *s* approaches 0 and expand Equation S9 to order *s*^4^.

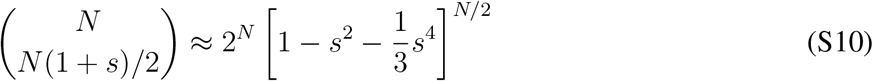

Next, we assume *N* is large and express the term under the brackets as an exponential.

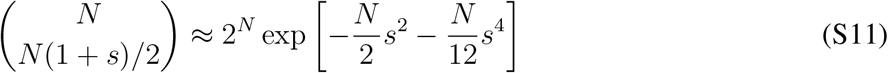

We can insert our approximate expression for the binomial coefficient back into *P* (*s*) (Equation S3) and obtain,

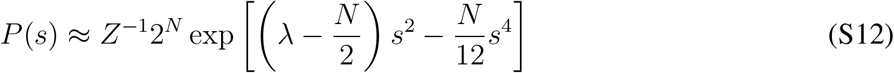

Note that Equation S12 (after transforming into free energy space) maps onto Landau theory (Equation S4). *s* corresponds to the order parameter and *λ*_*c*_ = *N/*2. At *λ* = *λ*_*c*_, *P* (*s*) switches from unimodal to bimodal, corresponding to a second order phase transition. We report a rescaled version of *λ* called Λ in Figure 5 and in other places in the Supporting Information to easily gauge how far an individual’s connection strength is from the critical point.

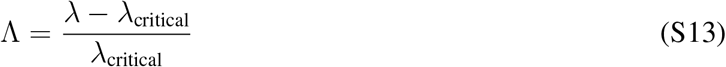

### Alternative N_eff_ fitting approach

Rather than choose one *N*_eff_ for all individuals in the data set as done in the main text, we could fit *N*_eff_ for each individual. Figure S1 demonstrates that such a procedure results in *N*_eff_ values that are highly linearly related with *λ*. In other words, more precise *N*_eff_ fits do not provide any more insight than maximum entropy fits of *λ* for all individuals in a data set under one optimal *N*_eff_.

**Figure S1.**
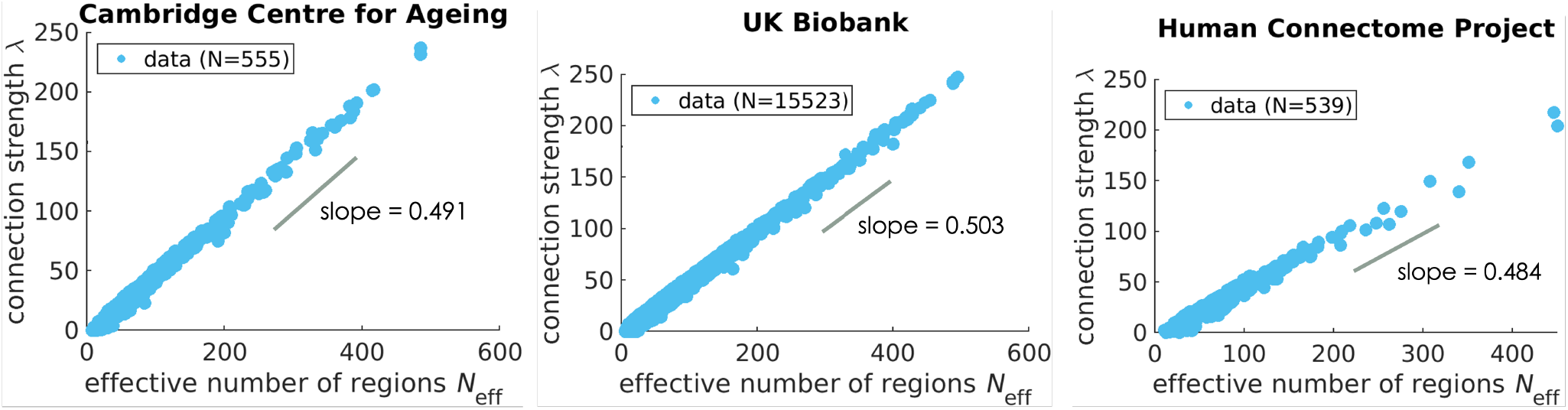
Treating *N*_eff_ as a parameter and fitting it per individual yields a strong correlation with *λ*. Each point reflects an individual brain scan and N reflects the total number analyzed. The value of the slope corresponds to that of the best-fit line for the data and is close to the predicted value of 0.5 (Equation S15). N is smaller than that of Figure 3 because some scans failed to have a minimum ⟨*s*^4^⟩ RMSE within the explored bounds of *N*_eff_ (4-500) or *λ* values were nonphysical by being less than 0.

The *N*_eff_-*λ* relationship can be reasoned from the analytical expression for *P* (*s*) (Equation S12). When Λ *<* 0, which many individuals satisfy (Figure S10), *P* (*s*) is well-approximated as a Gaussian.

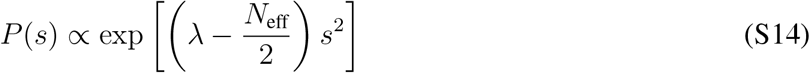

Thus, the analytical form for ⟨*s*^2^⟩ is:

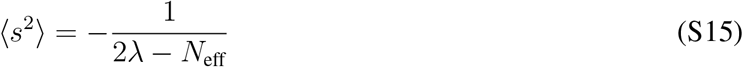

Since *λ* is fit in the Maximum Entropy framework to exactly match ⟨*s*^2^⟩, Equation S15 indicates that a larger *N*_eff_ requires a larger *λ* for a fixed ⟨*s*^2^⟩. Indeed, we find in Figure S1 that the best fit line of the *N*_eff_-*λ* relationship has an approximate slope of 0.5, in agreement with Equation S15.

### More supporting information

**Table S1.** Data set values for *P*_seg_ calculations under our particular fMRI preprocessing procedure (Methods).

| Data set | effective number of regions $N_{\text{eff}}$ | synchrony threshold $s^*$ |
| --- | --- | --- |
| Cambridge Centre for Ageing | 40 | 0.334 |
| UK Biobank | 30 | 0.357 |
| Human Connectome Project | 40 | 0.334 |

**Figure S2.**
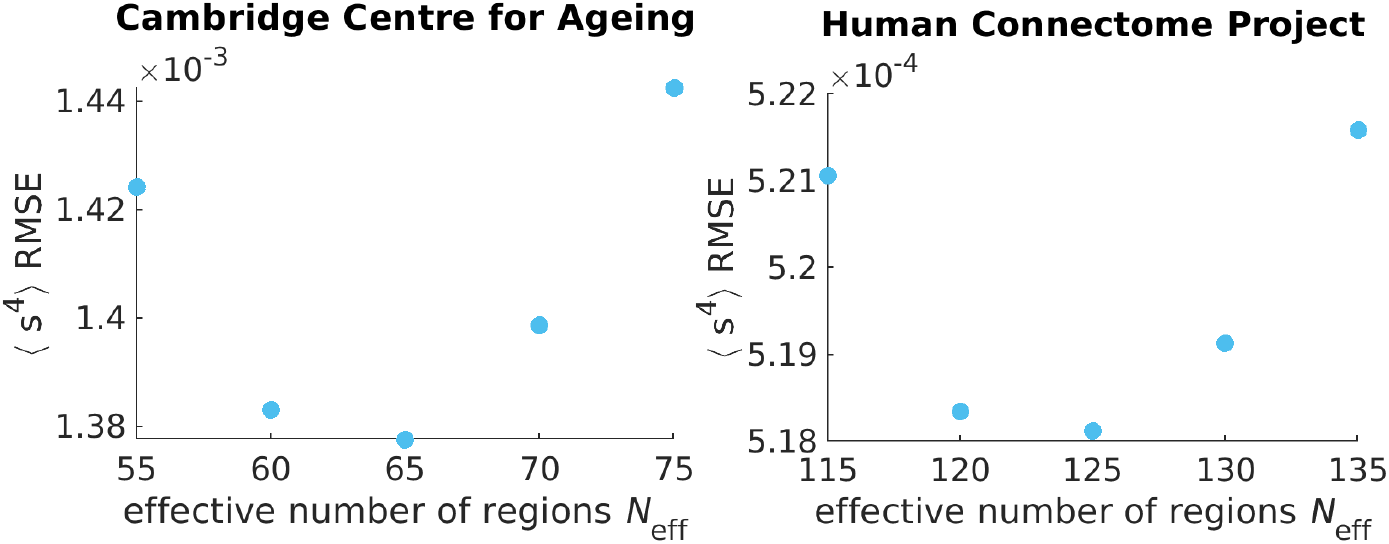
Identifying the effective number of regions *N*_eff_ for brain scans processed at the voxel-level. Each data point corresponds to the sum over all individuals’ RMSEs in the respective data set.

**Figure S3.**
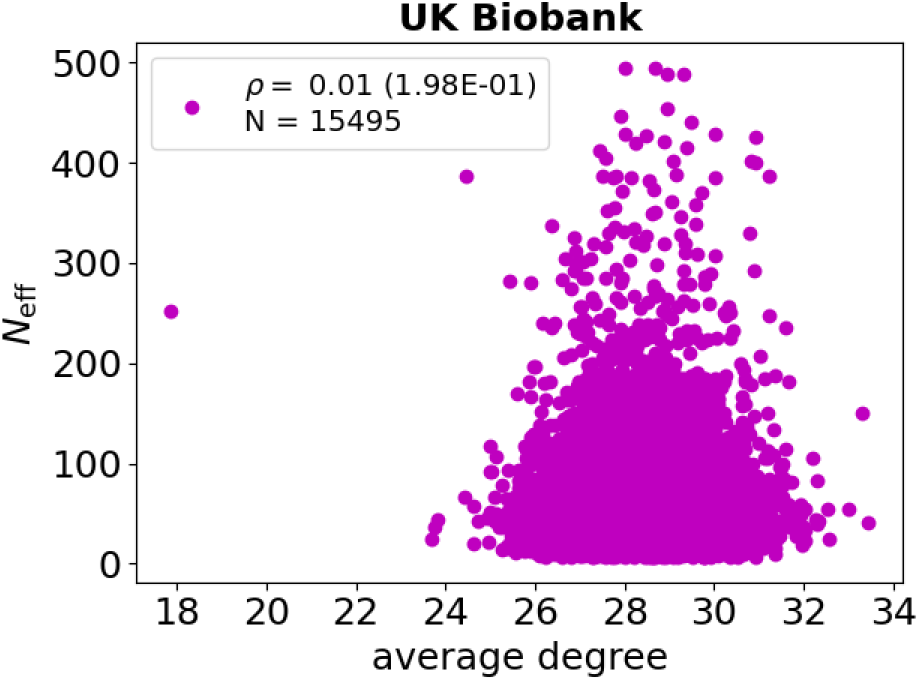
Individually fitted *N*_eff_ values from Figure S1 are not related to the average number of white matter tracts per brain region (average degree) as determined by diffusion MRI. The Q-Ball tractography method is used to analyze diffusion MRI scans (Methods). Data points correspond to individuals. The variable *ρ* corresponds to the Spearman correlation coefficient between average degree and *N*_eff_ calculated over all N individuals, with the p-value in parenthesis.

**Figure S4.**
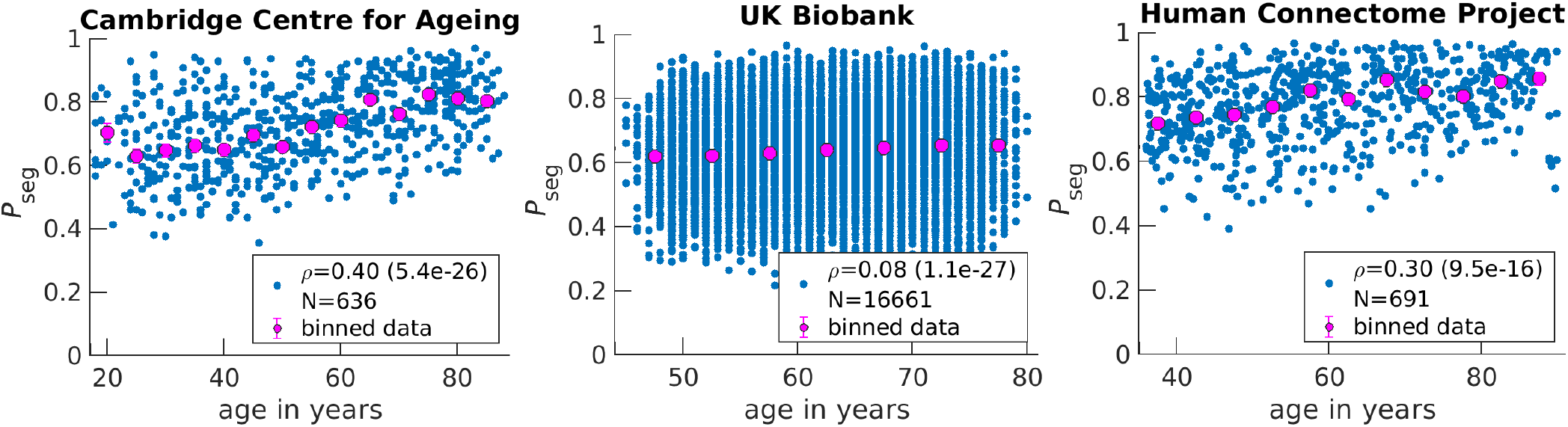
*P*_seg_ rises on average in aging brains but varies greatly among individuals with the same age. Blue data points correspond to individuals. The variable *ρ* corresponds to the Spearman correlation coefficient between age and *P*_seg_ calculated over all N individuals, with the p-value in parenthesis. Magenta points are the exact same data points presented in Figure 3 for the corresponding data set. Note that the corresponding error bars are not visible in these plots.

**Figure S5.**
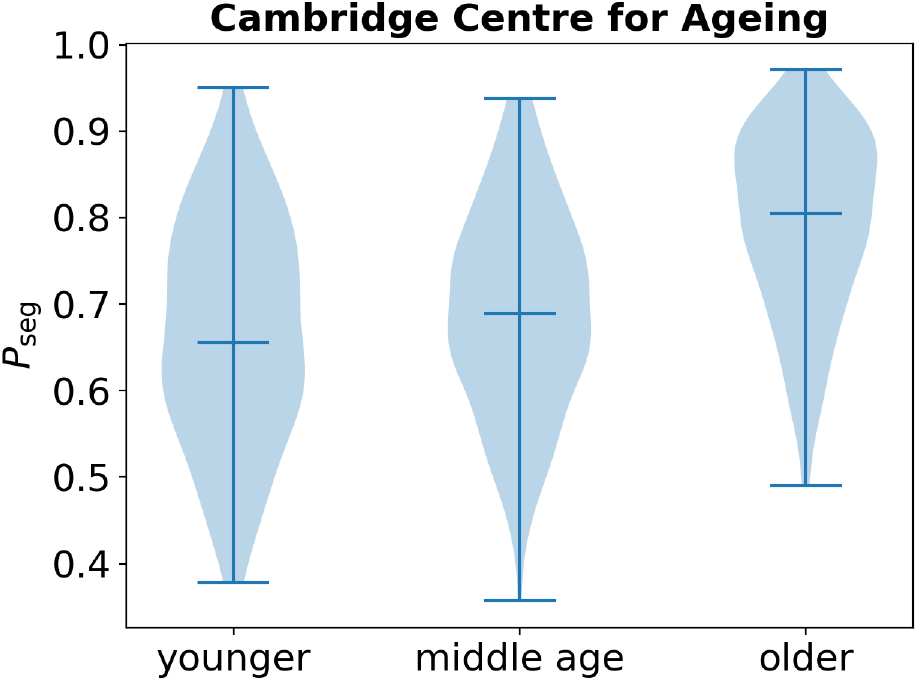
*P*_seg_ rises in aging brains across three Cambridge Centre for Ageing and Neuroscience age groups. Violin plots are presented, where middle horizontal lines correspond to medians while lower and upper lines correspond to minimum and maximum values, respectively. Younger individuals are those less than 35 years old (N=117); middle age, 40-60y (N=187); older, above 65y (N=209).

**Figure S6.**
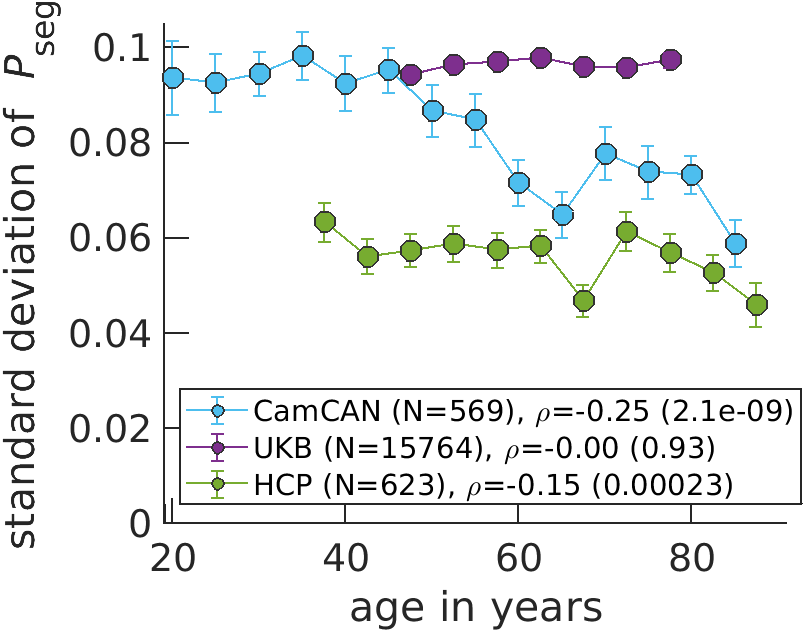
Standard deviations of *P*_seg_ per individual decreases as a function of age for CamCAN and HCP data sets. Data points correspond to medians, while error bars correspond to standard errors for bins of 5 years. The variable *ρ* corresponds to the Spearman correlation coefficient between age and *P*_seg_ calculated over all N individuals, with the p-value in parenthesis. Here, fMRI time-series data for an individual are equally split into 5 chunks and *P*_seg_ is calculated for each chunk before taking its standard deviation. In the main text, fMRI data are not split up and the entire time-series is considered in calculating *P*_seg_.

**Table S2.** Linear regression results for *P*_seg_ as a function of age

| CamCAN | coefficient | $t$ statistic | Prob> $\vert t\vert $ | UK Biobank | coefficient | $t$ statistic | Prob> $\vert t\vert $ |
| --- | --- | --- | --- | --- | --- | --- | --- |
| intercept | 0.575 | 40.0 | 8.66E-176 | intercept | 0.556 | 69.1 | <1E-300 |
| age | 0.0028 | 11.1 | 3.03E-26 | age | 0.0013 | 9.92 | 4.06E-23 |

| HCP | coefficient | $t$ statistic | Prob> $\vert t\vert $ |
| --- | --- | --- | --- |
| intercept | 0.650 | 38.1 | 1.08E-171 |
| age | 0.0021 | 7.58 | 1.14E-13 |

**Table S3.** Multiple linear regression results for *P*_seg_ as a function of age and sex across the data sets.

| CamCAN | coefficient | $t$ statistic | Prob> $\vert t\vert $ | UK Biobank | coefficient | $t$ statistic | Prob> $\vert t\vert $ |
| --- | --- | --- | --- | --- | --- | --- | --- |
| intercept | 0.583 | 39.0 | 1.22E-170 | intercept | 0.561 | 70.8 | <1E-300 |
| sex(T.male) | -0.0163 | -1.75 | 8.03E-02 | sex(T.male) | -0.0445 | -23.7 | 6.17E-122 |
| age | 0.0028 | 11.1 | 1.58E-26 | age | 0.0015 | 12.1 | 8.79E-34 |

| HCP | coefficient | $t$ statistic | Prob> $\vert t\vert $ |
| --- | --- | --- | --- |
| intercept | 0.668 | 39.6 | 1.92E-179 |
| sex(T.male) | -0.0500 | -6.17 | 1.20E-09 |
| age | 0.0022 | 8.03 | 4.21E-15 |

**Table S4.** Multiple linear regression results for *P*_seg_ as a function of age, sex and handedness for the Human Connectome Project.

| HCP | coefficient | $t$ statistic | Prob> $\vert t\vert $ |
| --- | --- | --- | --- |
| intercept | 0.677 | 32.8 | 5.75E-143 |
| sex(T.male) | -0.0504 | -6.20 | 9.72E-10 |
| handedness(T.right) | -0.0095 | -0.76 | 4.48E-01 |
| age | 0.0022 | 8.00 | 5.59E-15 |

**Figure S7.**
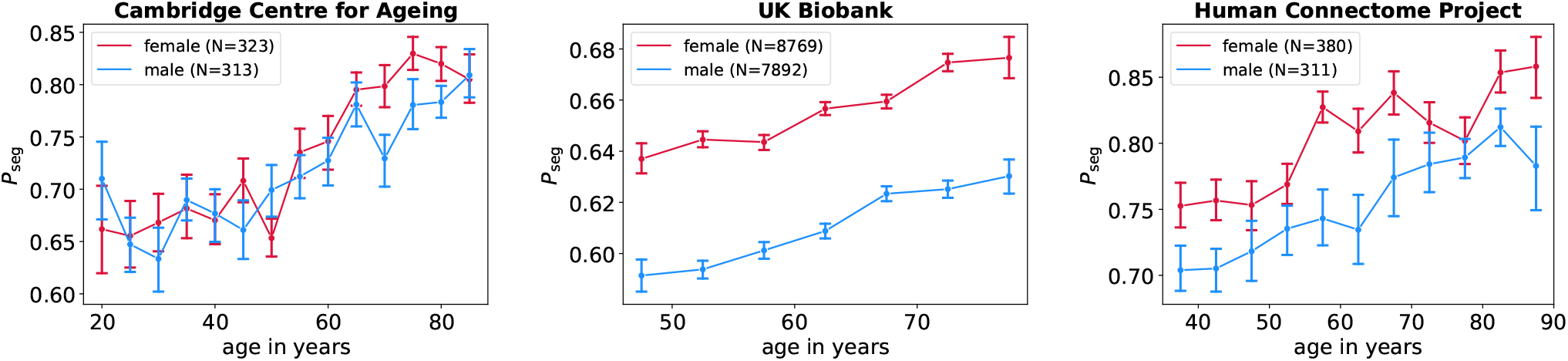
*P*_seg_ rises in aging brains across three data sets regardless of sex. Data points correspond to medians, while error bars correspond to standard errors for bins of 5 years. For UKB and HCP, we find that females’ brains have higher shifted *P*_seg_ values across age.

**Figure S8.**
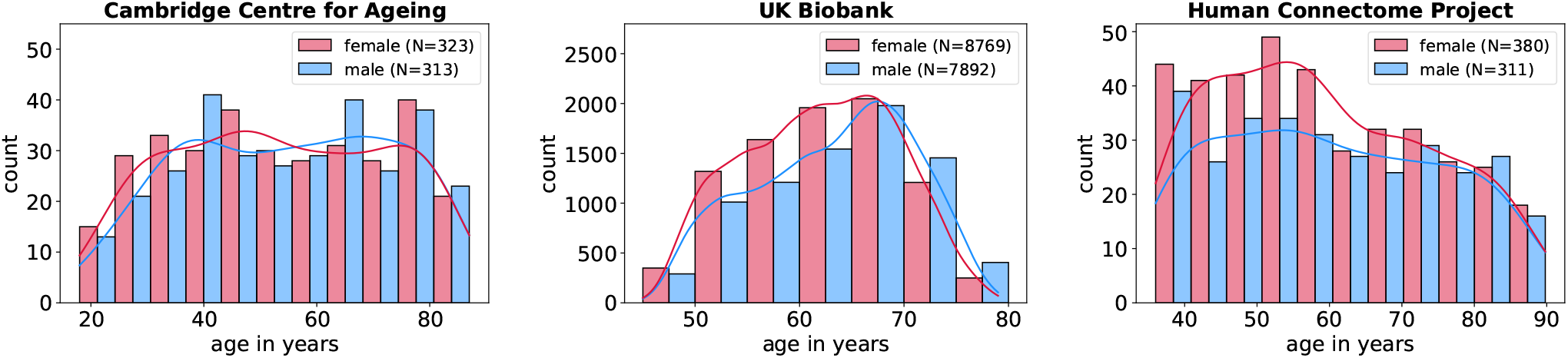
Sex is fairly well-represented across age across the three data sets. Thus, observed *P*_seg_ aging trends cannot be attributed to the increasing over-representation of one sex.

**Figure S9.**
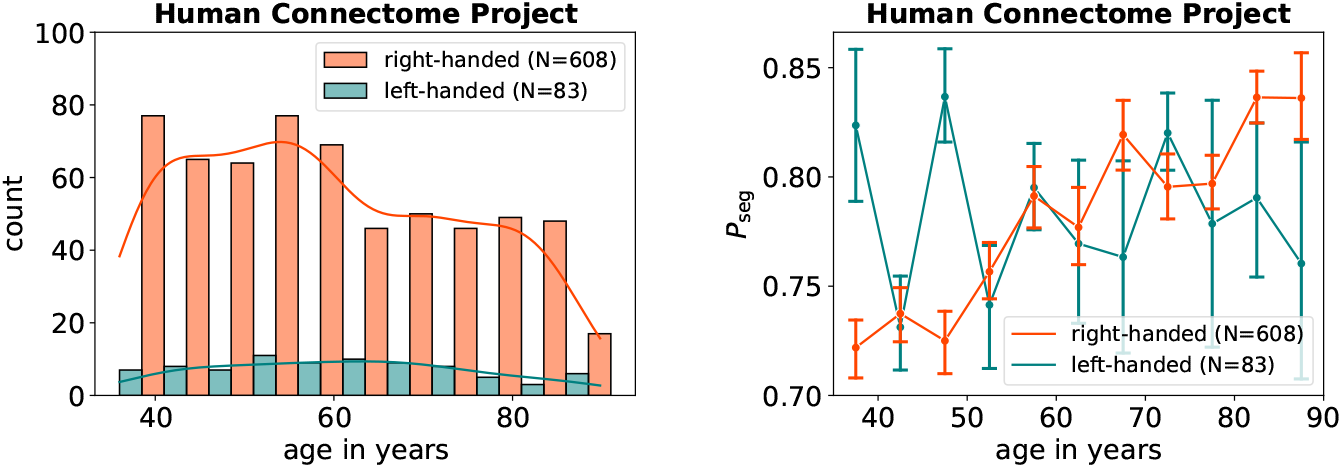
*P*_seg_ roughly rises in aging brains regardless of handedness. Data points correspond to medians, while error bars correspond to standard errors for bins of 5 years. Large error bars are seen for left-handed individuals because of small sample sizes (left plot).

**Figure S10.**
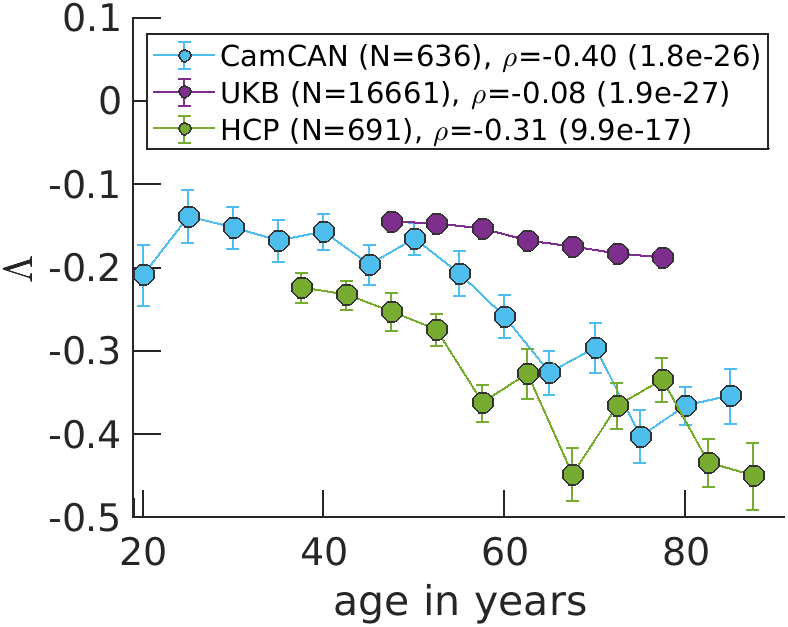
The rescaled connection strength parameter Λ moves further away from the critical point (Λ = 0) as age increases. Trends are similar in form to Figure 3 because *P*_seg_ is a function of Λ (Equation 2). Data points correspond to medians, while error bars correspond to standard errors for bins of 5 years. The variable *ρ* corresponds to the Spearman correlation coefficient between age and Λ calculated over all N individuals, with the p-value in parenthesis.

**Figure S11.**
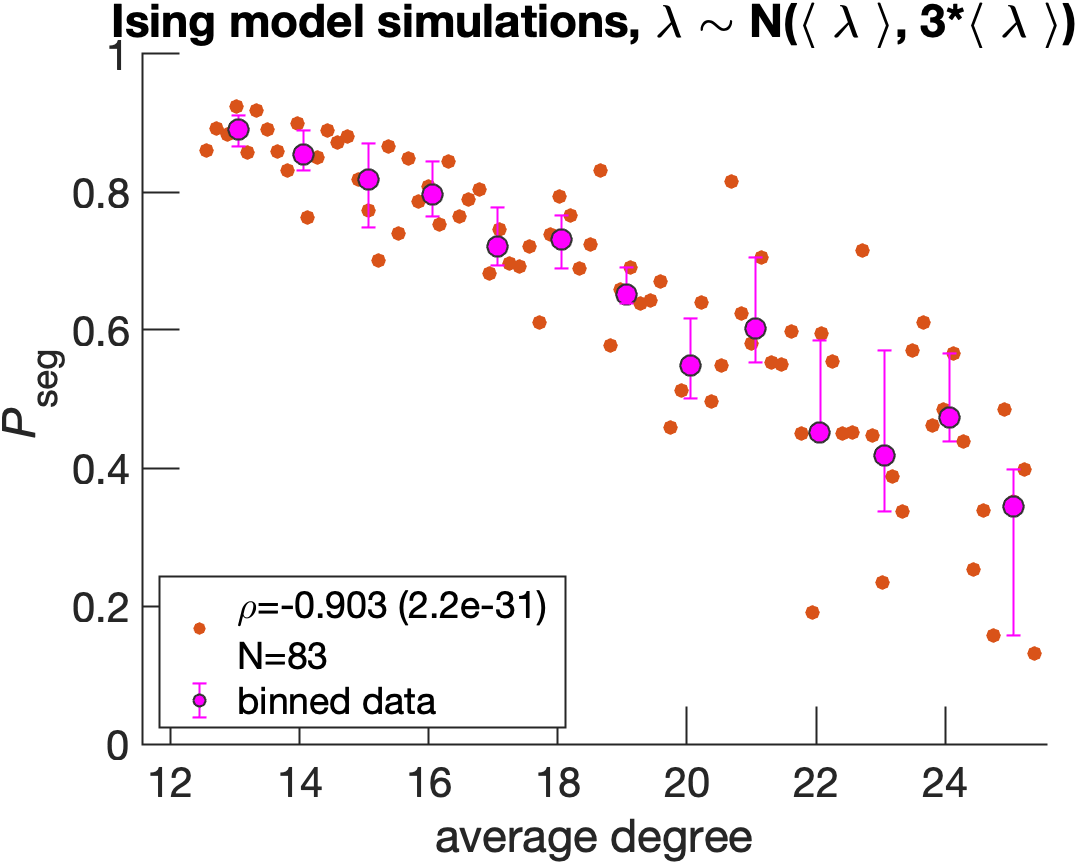
Greater variance in simulations is seen when edges’ connection strengths *λ* are drawn from a normal distribution with mean *(λ)* and standard deviation 3 * *(λ)*. At each consecutive step, *(λ)* is attenuated such that 5 edges are effectively removed per step (*(λ*^*1*^) = *(λ)p*_edge_) from the same starting dMRI structure as in Figure 4 (UK Biobank subject ID: 6025360). Data points correspond to medians, while error bars correspond to standard errors for bins of 5 years. Orange data points on the right plot correspond to individual Ising systems, where N reflects the total number. The variable *ρ* corresponds to the Spearman correlation coefficient calculated over all orange data points between average degree and *P*_seg_, with the p-value in parenthesis. Magenta data points correspond to medians, while error bars correspond to upper and lower quartiles for bin sizes of one degree.

**Figure S12.**
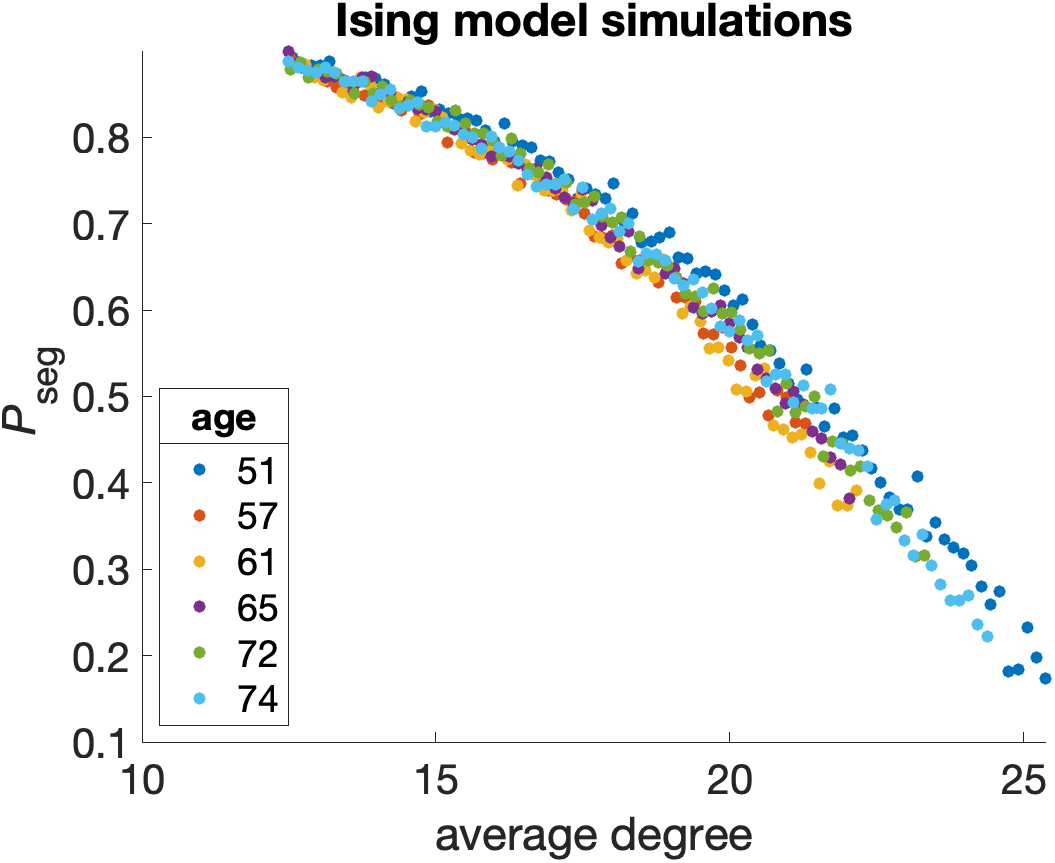
Similar results for Ising simulations are seen as in Figure 4 for different UK Biobank individuals with different ages. Edges are randomly removed as in Figure 4. Starting diffusion MRI structures are used from following subject IDs: 6025360 (51y), 4712851 (57y), 3081886 (61y), 1471888 (65y), 4380337 (72y), and 1003054 (74y).

**Figure S13.**
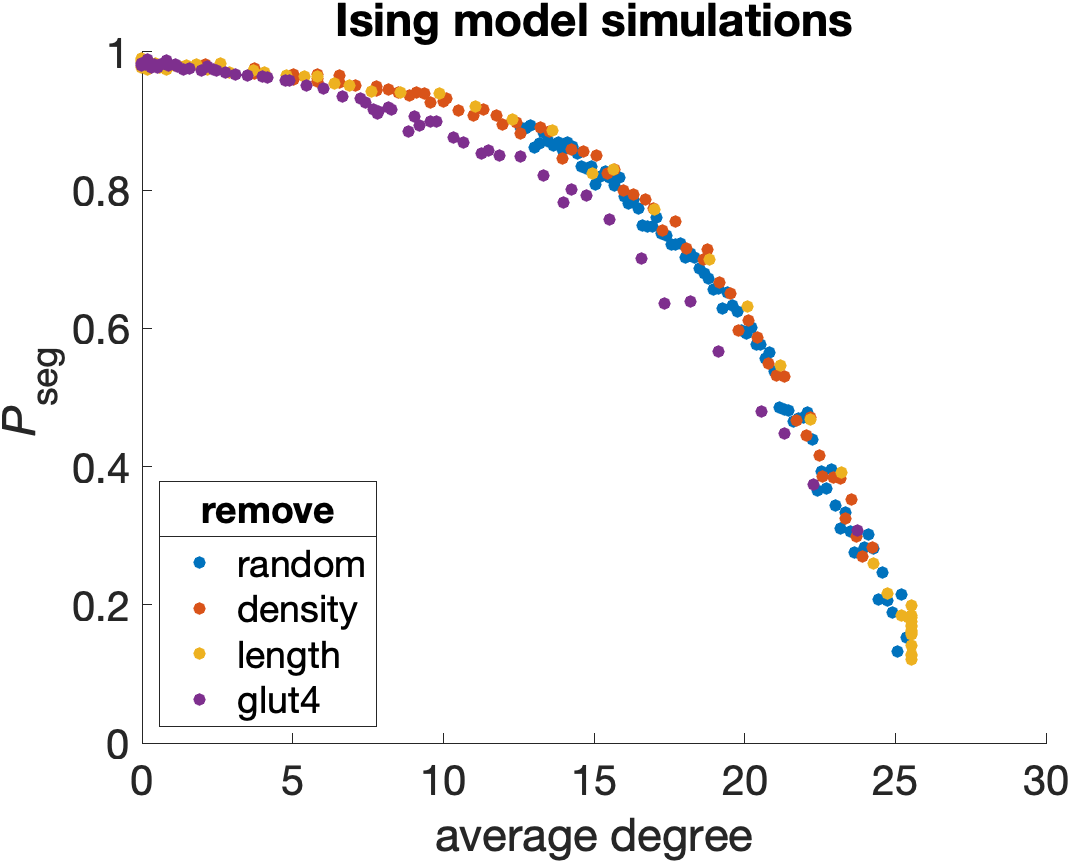
Edge removal mechanisms only matter in so much as they attenuate average degree for Ising simulations. In addition to randomly removing edges as shown in Figure 4, we computationally remove edges based on targeted attack of tract density, tract length, and a node’s GLUT4 receptor density. Edges are removed in sequential order, such that those with the largest value are removed first. For all properties except for random, we remove edges until none are present for the same starting dMRI structure as in Figure 4 (UK Biobank subject ID: 6025360).

**Figure S14.**
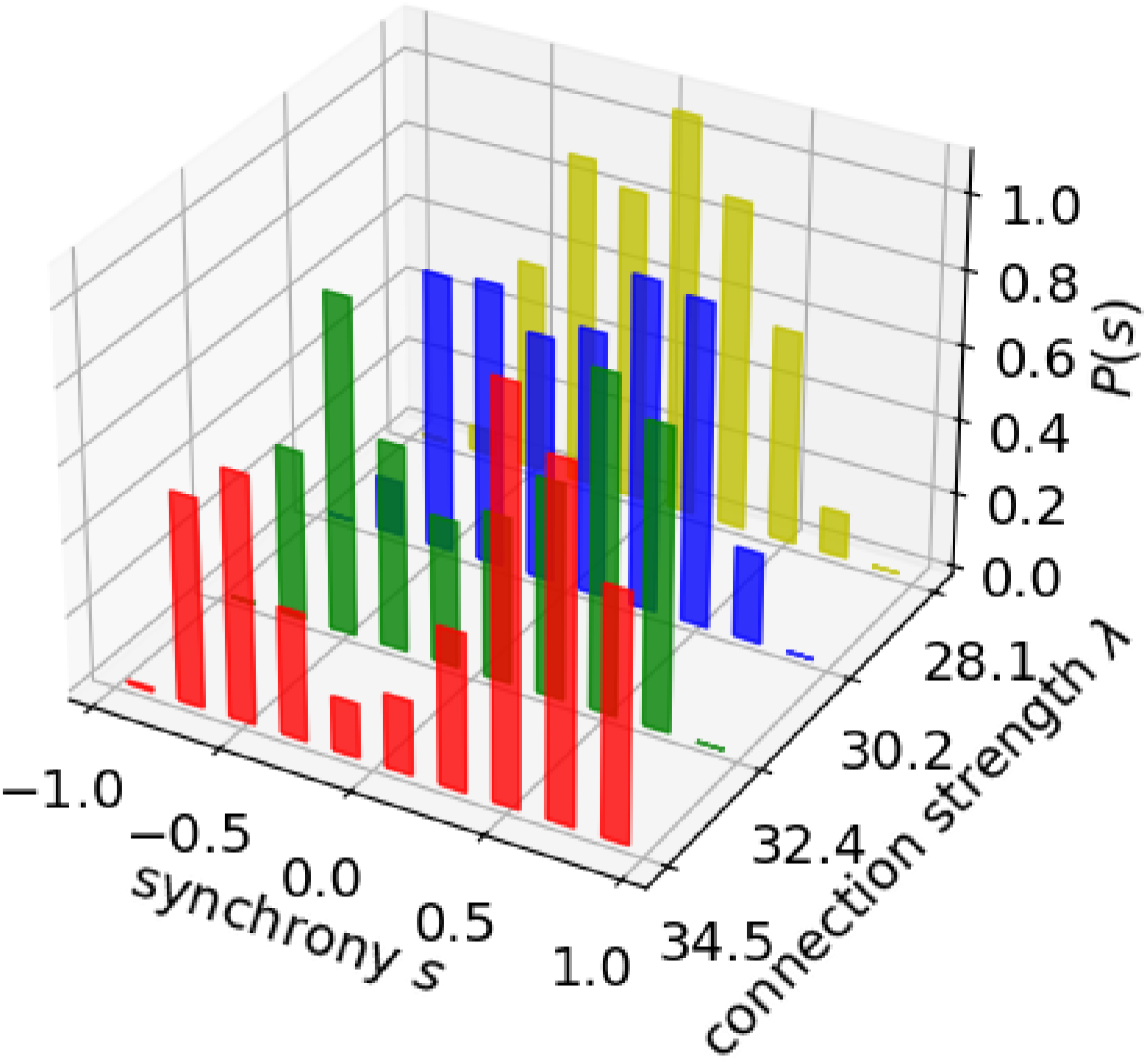
Synchrony distributions transform from bimodal to unimodal as edges are randomly removed from UK Biobank subject ID: 6025360. The parameter *λ* relates to edge removal because *λ* = *λ*_0_ * *p*_edge_, where *λ*_0_ is a constant throughout the edge removal process and *p*_edge_ is the probability that two nodes share an edge (Methods).

**Figure S15.**
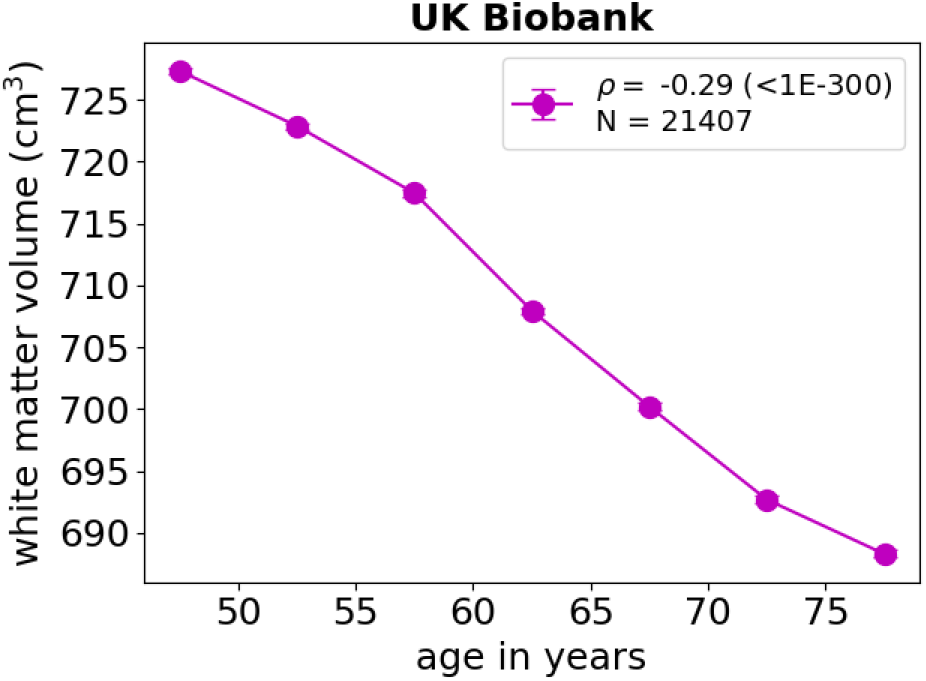
White matter volume decreases with age. White matter volume is measured by structural MRI provided by the UK Biobank. Data points correspond to medians, while error bars correspond to standard errors for bins of 5 years. The variable *ρ* corresponds to the Spearman correlation coefficient between age and white matter volume calculated over all N individuals, with the p-value in parenthesis. Error bars are plotted but are not visible because of their minuscule size. N is larger than that of Figure 3 because all individuals with structural MRI scans are considered.

**Figure S16.**
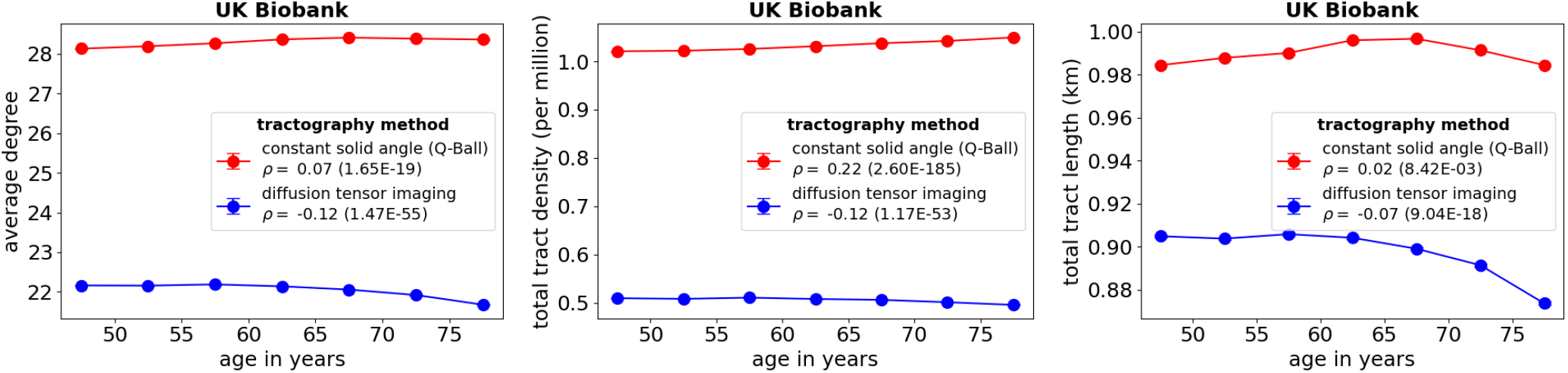
White matter tract properties do not degrade as a function of age when using the Q-Ball method for tractography. However, they do degrade with age when using the less accurate diffusion tensor imaging method. Data points correspond to medians, while error bars correspond to standard errors for bins of 5 years. The variable *ρ* corresponds to the Spearman correlation coefficient between age and the corresponding property calculated over all available individuals (N=16,649), with the p-value in parenthesis. Error bars are plotted but are not visible because of their minuscule size.

**Figure S17.**
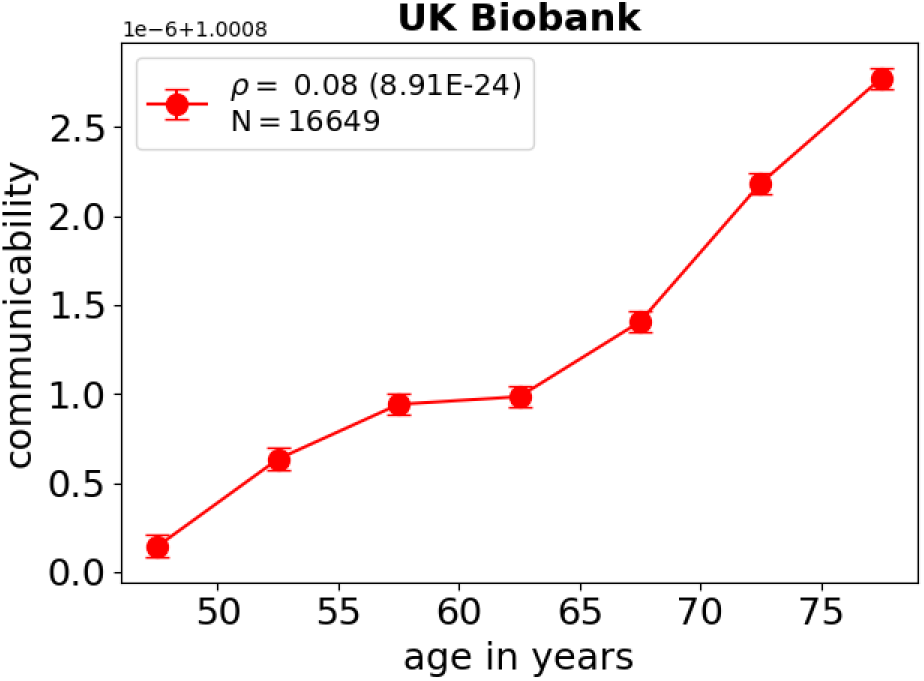
Mean communicability across all brain region pairs does not decrease with age. Communicability is calculated based on tract density as measured by the Q-Ball method for tractography (Methods). Data points correspond to medians, while error bars correspond to standard errors for bins of 5 years. The variable *ρ* corresponds to the Spearman correlation coefficient between age and white matter volume calculated over all N individuals, with the p-value in parenthesis. Note that the y-axis should be scaled by 10^−6^ and shifted by 1.0008.

**Figure S18.**
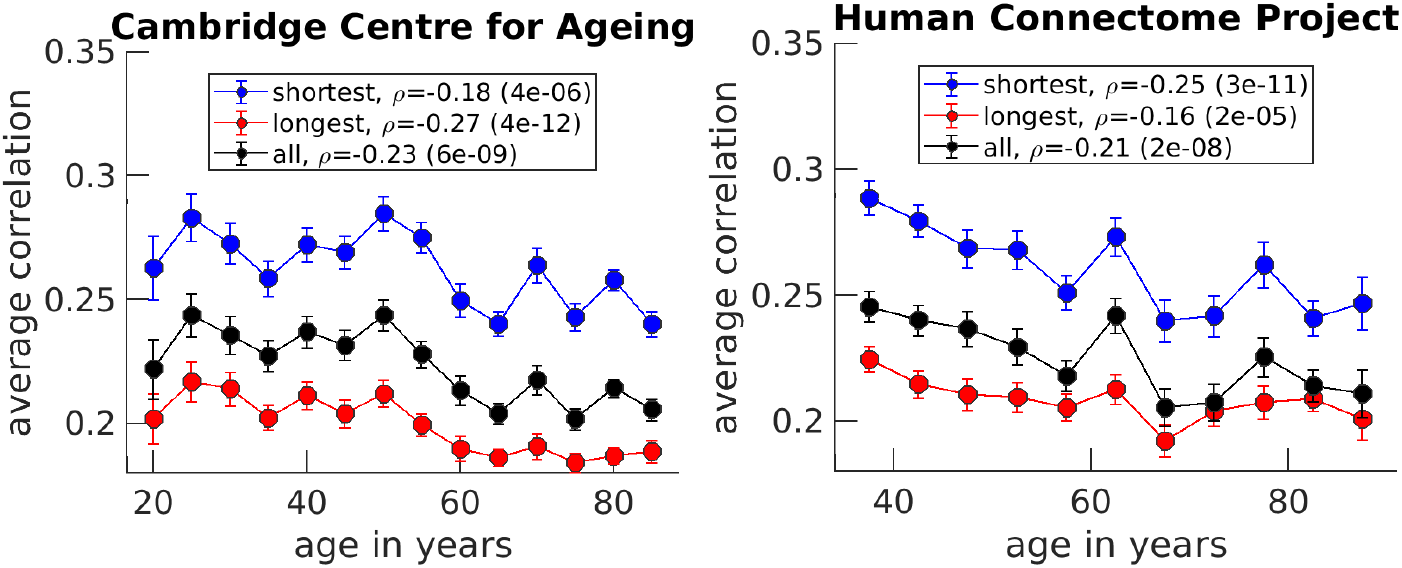
For the Cambridge Centre for Ageing and Neuroscience data set, the shortest edges (lower quartile) have average Pearson correlations or average functional connectivities which correlate less than those of the longest edges (upper quartile). For the Human Connectome Project, the opposite is the case. Edge distances are measured by center of mass coordinates of the brain regions based on the Seitzman atlas. Shortest and longest edges correspond to the lower and upper quartile (25%), respectively. Only positive correlations are considered and diagonal elements are ignored. Data points correspond to medians, while error bars correspond to standard errors for bins of 5 years. The variable *ρ* corresponds to the Spearman correlation coefficient between age and average correlation calculated over all available individuals (N_CamCAN_ = 640 and N_HCP_ = 700), with the p-value in parenthesis.

**Table S5.** Functional MRI acquisition parameters of the data sets.

| Data set | field strength | repetition time | echo time | flip angle | voxel size | total time points |
| --- | --- | --- | --- | --- | --- | --- |
| CamCAN | 3T | 1970 ms | 30 ms | 78° | 3x3x4.44 mm<sup>3</sup> | 241 |
| UK Biobank | 3T | 735 ms | 39 ms | 52° | 2.4x2.4x2.4 mm<sup>3</sup> | 490 |
| HCP | 3T | 800 ms | 37 ms | 52° | 2x2x2 mm<sup>3</sup> | 1912 |

**Table S6.** Demographic information of the data sets for those individuals in Figure 3.

| Data set | age range | $\langle \text{age} \rangle \pm \text{std}(\text{age})$ | sex |
| --- | --- | --- | --- |
| CamCAN | 18-87 | 54.2±18.6 | 323F/313M |
| UK Biobank | 45-79 | 54.8±7.4 | 8769F/7892M |
| HCP | 36-90 | 59.6±14.9 | 380F/311M |

**Figure S19.**
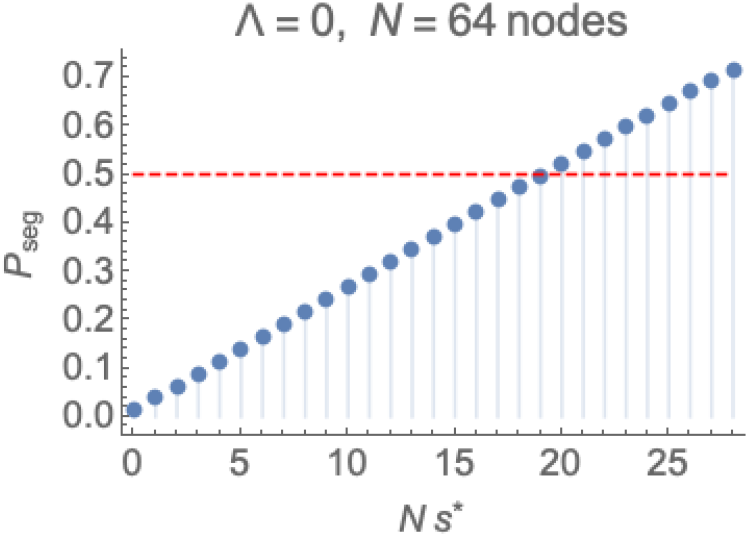
The synchrony threshold *s*^*^ is chosen such that it delineates between integrated and segregated states when *P*_seg_ = *P*_int_ = 1*/*2 (red line) at the critical point (Λ = 0). This particular figure is created for 64 nodes; it must be set to the corresponding data set’s *N*_eff_ to determine the appropriate synchrony threshold.

## TECHNICAL TERMS

**Integration** a network state composed of global signaling.

**Segregation** a network state limited to local signaling.

**State** a particular combination of physical properties. Here, we assume that brain networks can only occupy either the integrated or segregated state.

**Ising model** a classic model in physics that was first applied to ferromagnetism. It includes pairwise interactions between binary spin states.

**Phase** interchangeable with the word ‘state’ for the purposes of this text.

**Critical Point** the point where two phases coexist. In this text, it is where the synchrony distribution dramatically changes from bimodal (primarily integrated) to unimodal (primarily segregated).

**Maximum Entropy fit** a fitting strategy that satisfies user-defined constraints in the most agnostic way.

**White Matter** bundles of axons connecting brain regions.

